## Supplementary figures and tables for "Feature-dependent decorrelation of sound representations across the auditory pathway"

**Supplementary figures, Gosselin et al.**

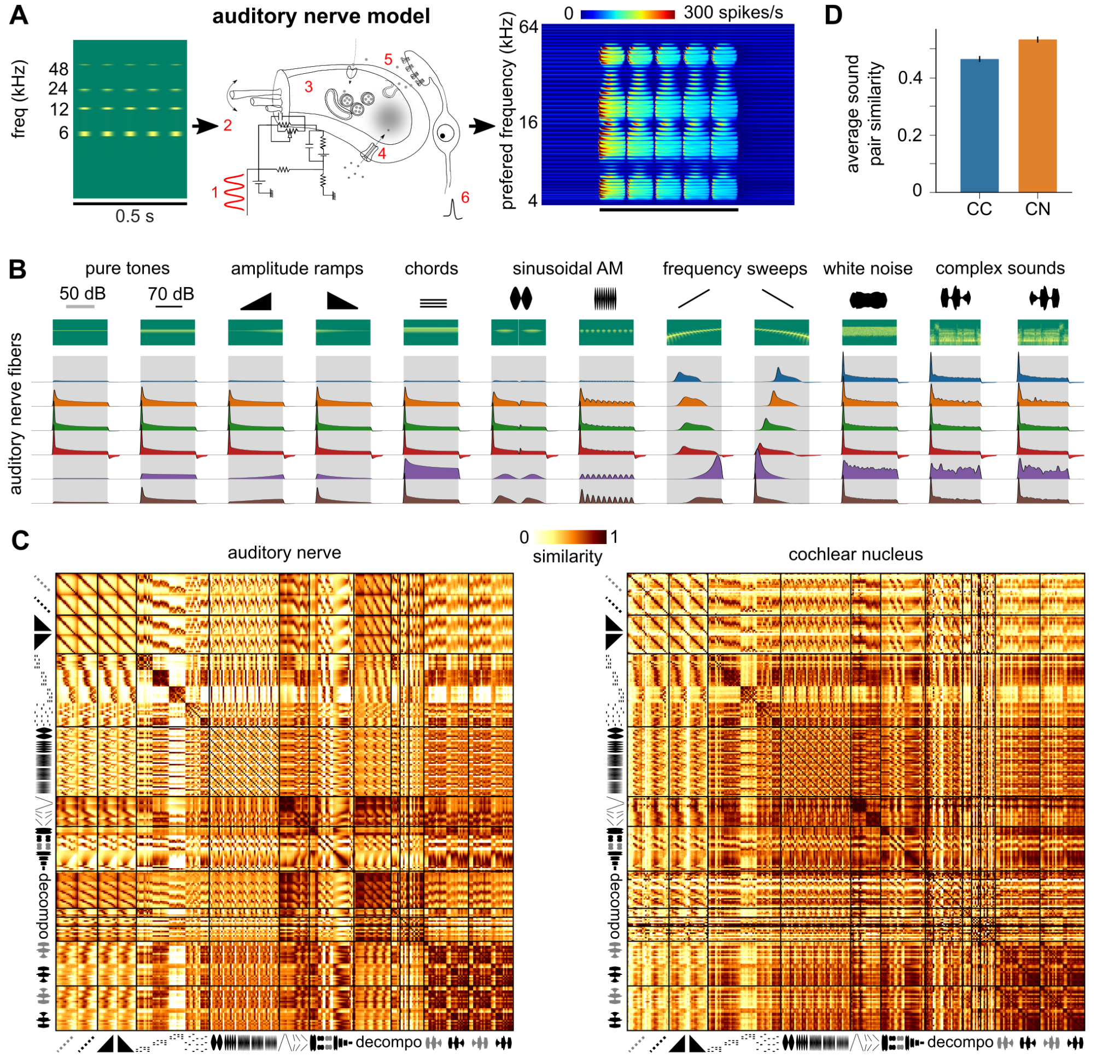

**Figure S1**: **Comparison of representations between cochlear nucleus and auditory nerve fiber model.** **A.** Sounds sampled at 400 kHz (represented here as a spectrogram) are fed into a detailed biophysical model of the cochlea and auditory nerve. The model simulates passive basilar membrane properties (1), stereocilia transduction (2, 3), calcium channel dynamics in hair cells (4) and synaptic release, as summarized in the Methods section. **B.** Responses of example neurons from the auditory nerve fiber model with spectral content at 12 kHz (2 pure tones, 2 ramps, 1 chord, 2 AMs, 2 chirps, 1 WN and 2 complex, represented with their spectrograms). Sound presentation periods are shaded in gray. **C.** Matrices of spatial representation similarity for the auditory nerve fiber model activity during sound presentation and for cochlear nucleus data as in **Fig. 3**. **D.** Average spatial representation similarity between all pairs of sounds computed from **C** for the auditory nerve fiber model and for cochlear nucleus data.

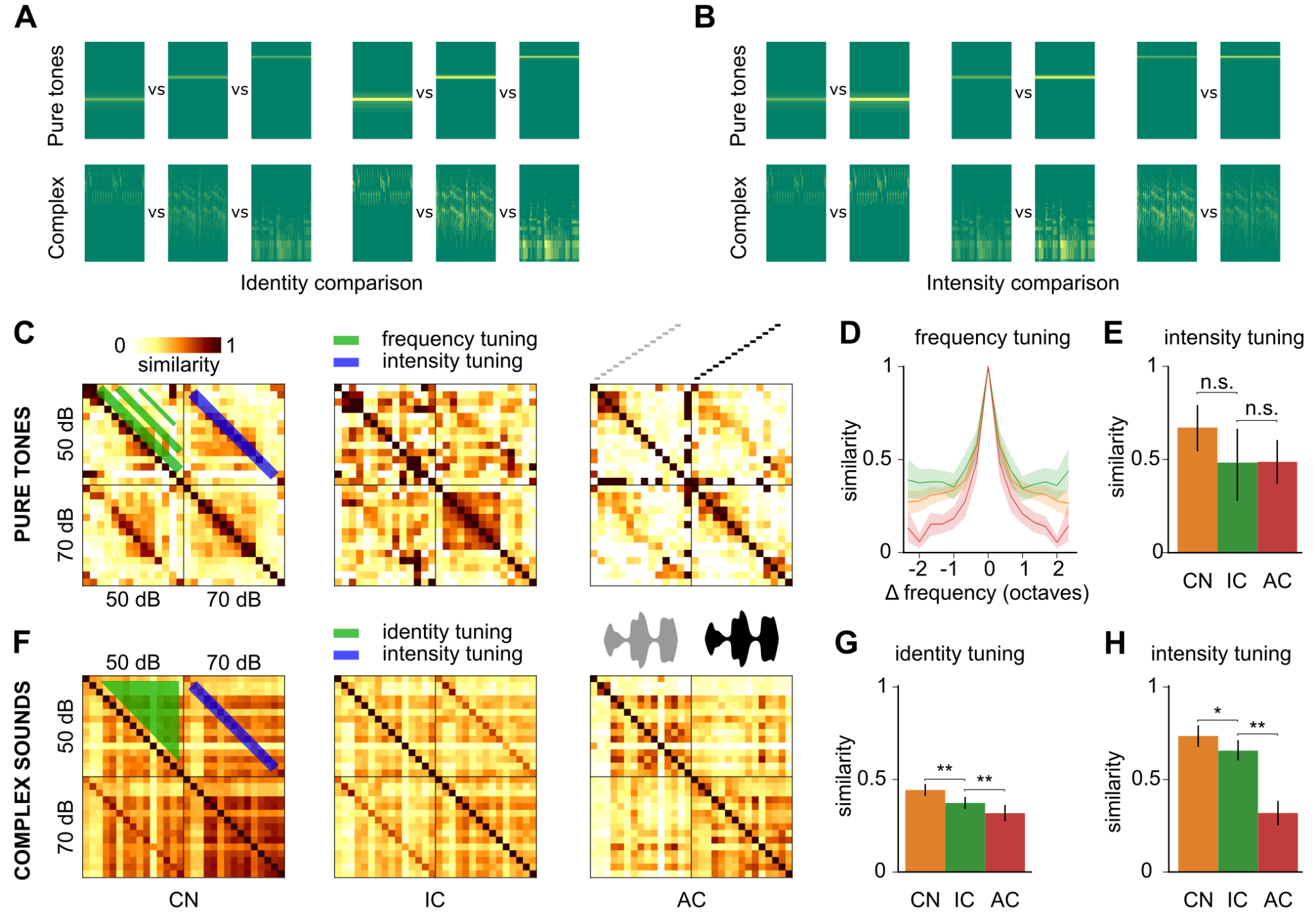

**Figure S2**: **Representation similarity of sound identity and intensity when including temporal information.** **A & B.** Sample spectrograms of the sounds compared in the population tuning analyses presented in **Fig. 4** and **S2C-H** separating identity and intensity comparisons. **C-H** Same measures of population tuning as in **Fig. 4A-F** but taking in account temporal information by comparing the concatenated population vector time series describing the full spatio-temporal population representation over the duration of the sound instead of using only the time-averaged activity. Temporal information does not change representation similarity for pure tones and pure tone intensity (**C-E**), but decreases similarity for complex sound identity and intensity (**F-H**). For identity, the result is expected as complex sounds include a large amount of temporal information. Complex sound decorrelation is therefore less pronounced for the spatio-temporal than for the time-averaged spatial representation (**Fig. 4E**), in line with the observation that a key computation of the auditory cortex is to encode temporal information into specific spatial patterns [[19]](https://www.zotero.org/google-docs/?Btclfv). The improvement of intensity tuning for complex sounds (**H**) and not for pure tones after including temporal information was also observed previously [[19]](https://www.zotero.org/google-docs/?x7NbUD). CN = cochlear nucleus, IC = inferior colliculus, AC = auditory cortex. Full statistics are provided in **Table S5**.

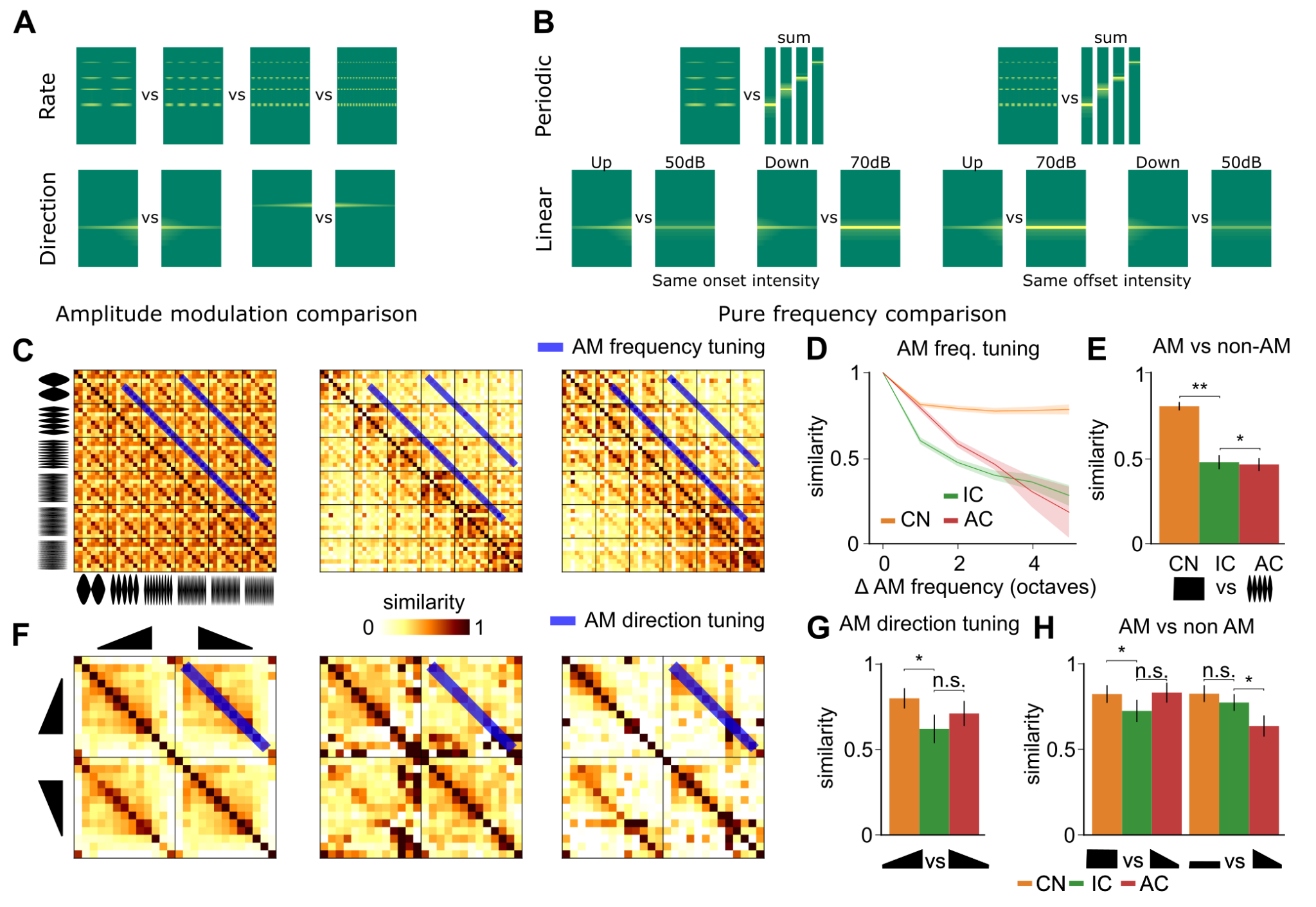

**Figure S3: Representation similarity of amplitude modulations when including temporal information. A.** Spectrogram of amplitude modulated sounds. **B.** Spectrogram summarizing the comparison between AM and non-AM activity. **C.** Spatio-temporal representation similarity matrices for 48 AM sounds at 8 carrier frequency contents and 6 modulation frequencies. **D.** Evolution of spatio-temporal similarity between AMs dependent on their modulation frequency difference. **E.** Spatio-temporal representation similarity between AM sounds and the non-modulated carrier signal. **F.** Spatio-temporal representation similarity matrices for upward and downward linear intensity ramps. **G.** Spatio-temporal similarity between upward and downward Ramps at the same frequency. **H.** Similarity between intensity ramps and pure tones at the same frequency and at the start (left) or end (right) intensity of the ramp. CN = cochlear nucleus, IC = inferior colliculus, AC = auditory cortex. Full statistics are provided in **Table S6**.

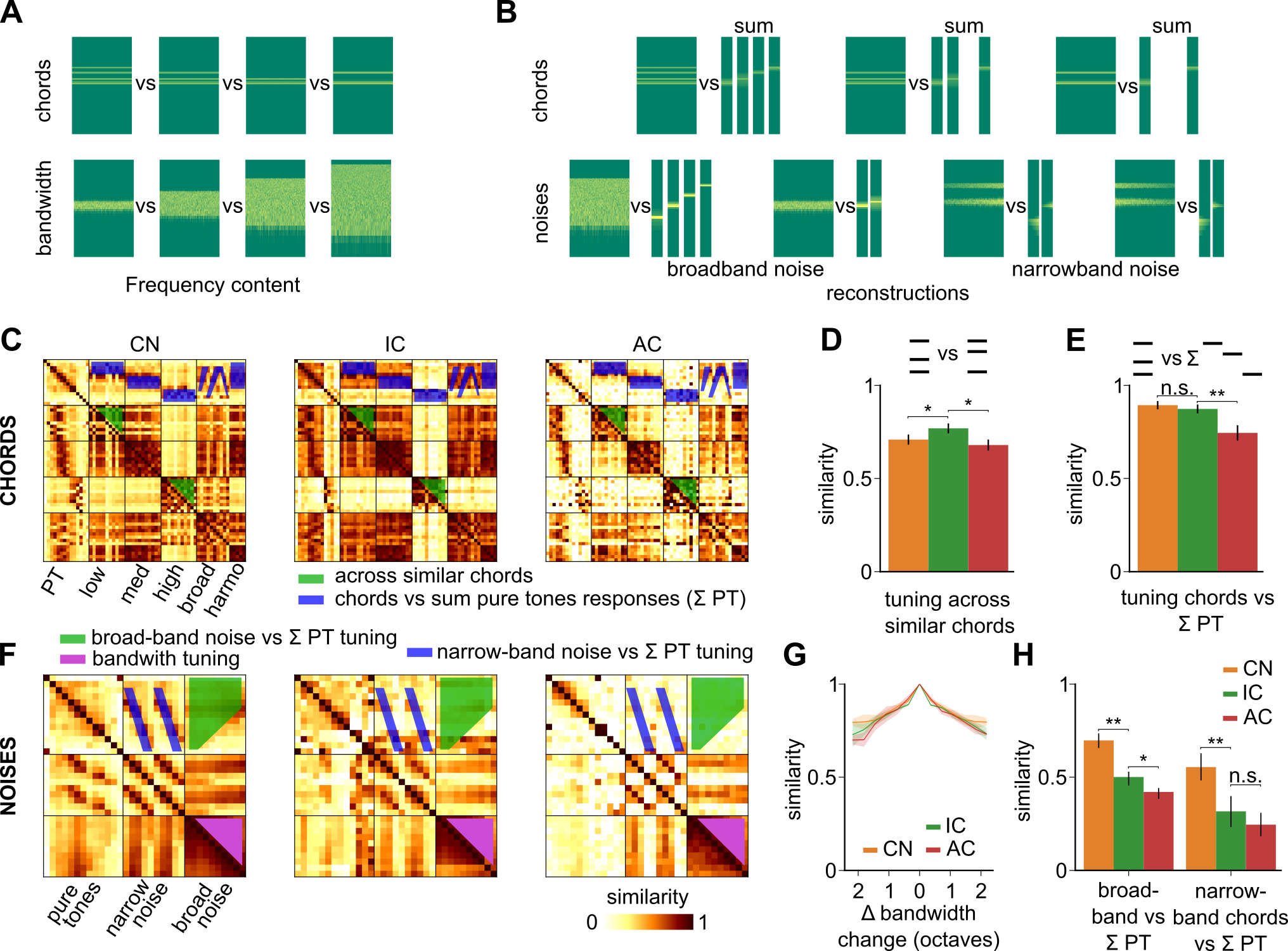

**Figure S4**: **Representation similarity of broad-band and multi-frequency sounds when including temporal information.** **A.** Spectrogram of different chords generated from the same pool of pure tones and of noises with different bandwidths. **B.** Spectrogram sketching the comparison between representation of 3 chords and the summed representations of the corresponding pure tones (top). Spectrogram sketching the comparison between representations of 4 noises and the summed representations of the pure tones included in their frequency range (bottom). **C.** Spatio-temporal representation similarity matrices of pure tones at 70 dB SPL and their summation into chords, organized based on their frequency content. **D.** Similarity between chords built from the same pool of pure tones. **E.** Similarity between spatio-temporal representations of chords and the summed representations of the pure tones that compose them. **F.** Spatio-temporal representation similarity matrices of pure tones, narrow- and broad-band noises. **G.** Spatio-temporal representation similarity between broadband noises dependent on their difference in bandwidth across recorded areas. **H.** Similarity of spatio-temporal representations between broadband (left) or narrowband (right) noises and the summed response of pure tones included in their frequency range. CN = cochlear nucleus, IC = inferior colliculus, AC = auditory cortex. Full statistics are provided in **Table S7**.

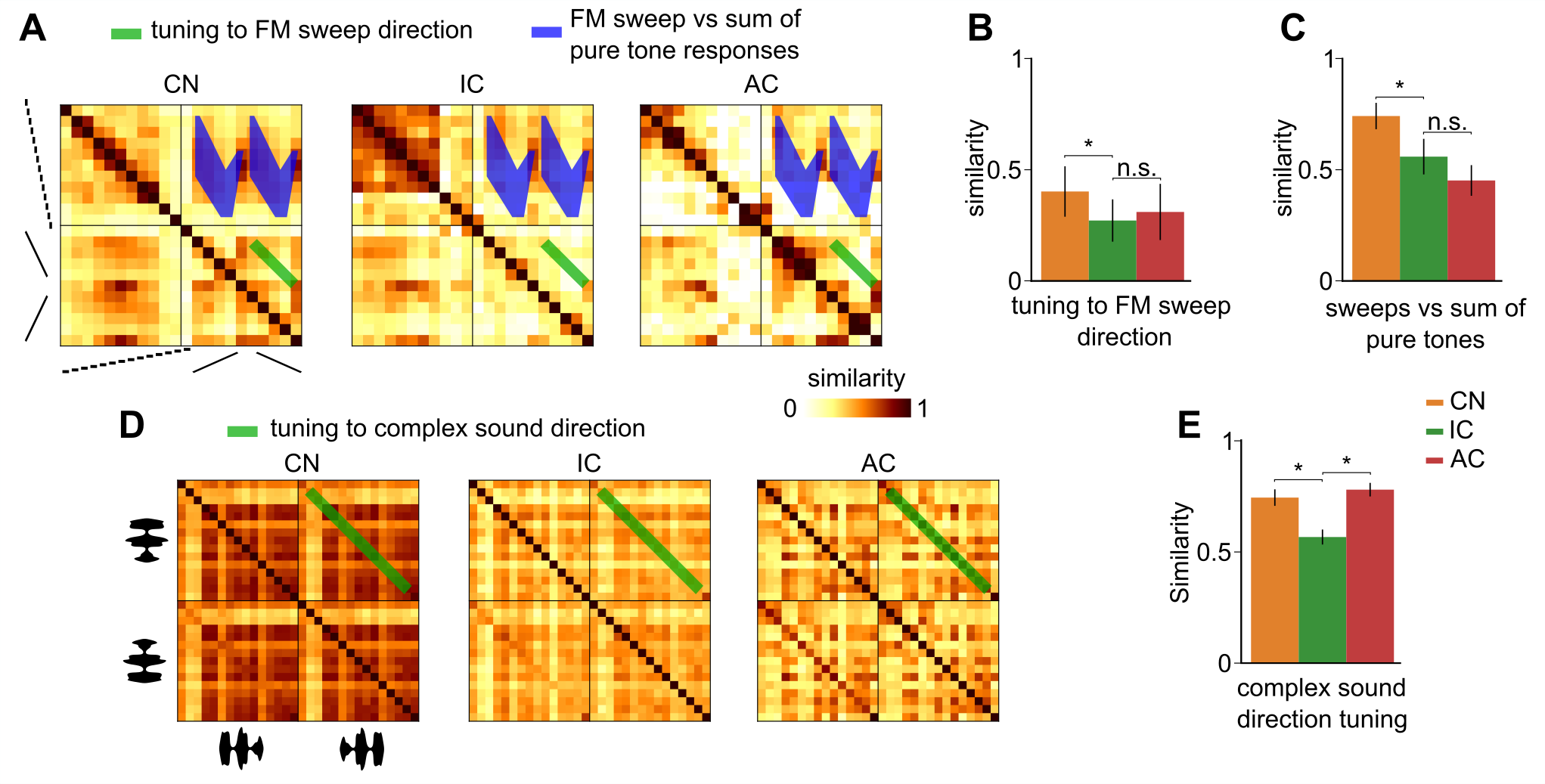

**Figure S5: Representation similarity for sound direction when including temporal information.** **A.** Representational similarity matrices of pure tones at 70 dB SPL and Chirps at 70 dB SPL that span the equivalent frequency range for the full spatio-temporal description of population activity in all regions. **B.** Similarity between same frequency up- and down-frequency modulated chirps for spatio-temporal population activity. **C.** Similarity between Chirps and the response reconstructed from pure tones traversed by the Chirp, for the spatial code. **D.** Representational similarity matrices of forward and backward Complex sounds at 70 dB SPL for spatio-temporal population activity in all regions. **E.** Similarity between the same complex sounds in both temporal directions for spatio-temporal population activity. CN = cochlear nucleus, IC = inferior colliculus, AC = auditory cortex. CN = cochlear nucleus, IC = inferior colliculus, AC = auditory cortex. Full statistics are provided in **Table S8**.

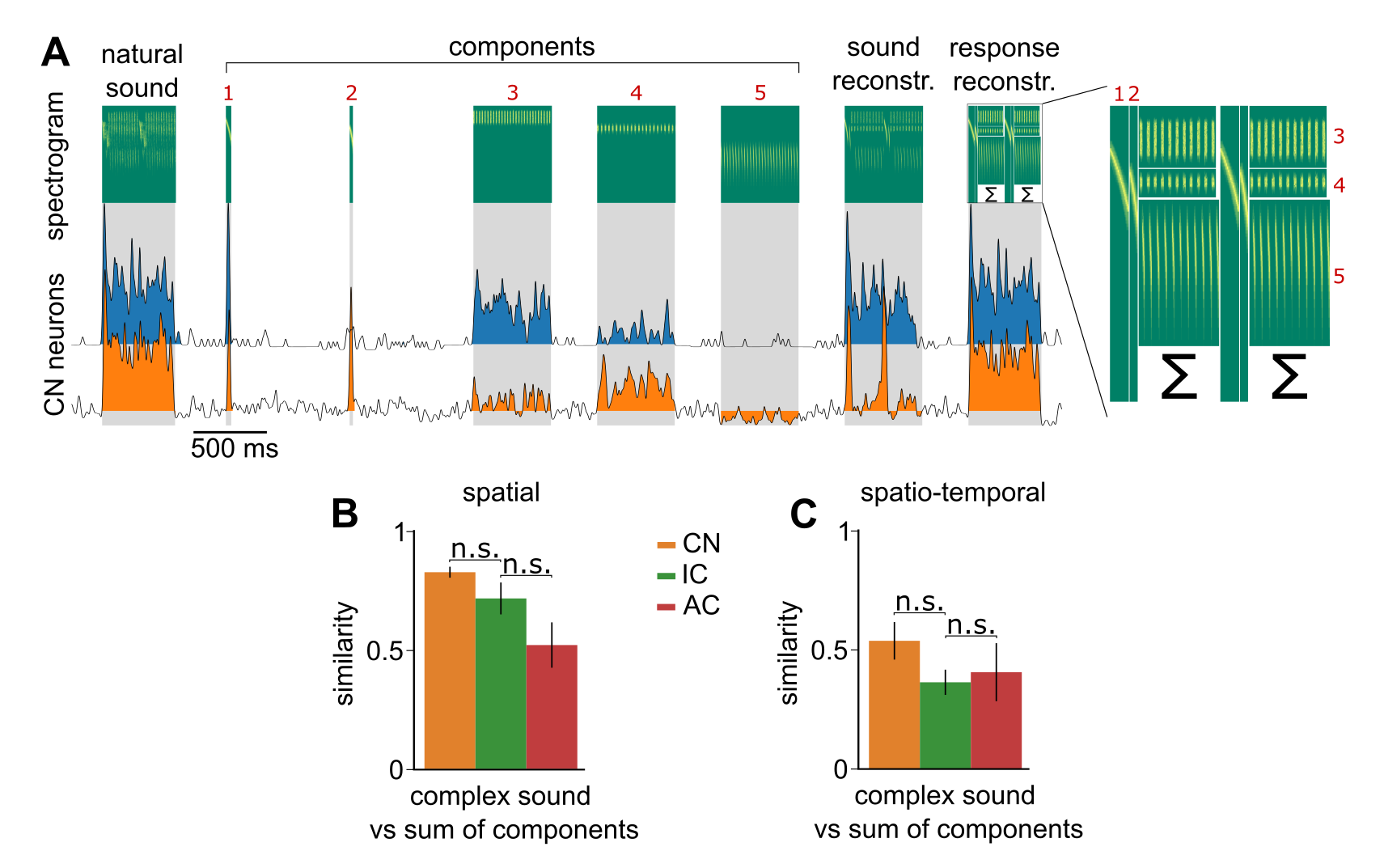

**Figure S6**: **Specificity of complex sound representations against the sum of their decomposition.** **A.** Schematic of the reconstruction process of sounds as a sum of individual components (top) and responses of 2 example neurons from the cochlear nucleus to the example complex sound, its components, its reconstruction, and its reconstructed response). **B.** Similarity between the responses to reconstructed complex sounds and reconstructed response from responses to components of the complex sound for 3 complex sounds for both codes. **C**. same as **B** but for the full spatio-temporal population activity. CN = cochlear nucleus, IC = inferior colliculus, AC = auditory cortex. Full statistics are provided in **Table S8**.

| **Identity coding** | | | | |
| --- | --- | --- | --- | --- |
| **Category** | **ΔOctaves** | **CN** | **IC** | **AC** |
| Pure tones | 0,3 | 0.62±0.05 | 0.57±0.09 | 0.52±0.08 |
|  |  | / | 8,59E-01 | 6,78E-01 |
|  | 0,6 | 0.43±0.03 | 0.46±0.05 | 0.32±0.08 |
|  |  | **/** | 4,01E-01 | 9,29E-02 |
|  | 0,9 | 0.38±0.03 | 0.33±0.05 | 0.22±0.06 |
|  |  | **/** | 3,98E-01 | 1,76E-01 |
|  | 1,2 | 0.35±0.02 | 0.34±0.06 | 0.18±0.04 |
|  |  | **/** | 9,17E-01 | 1,16E-01 |
|  | 1,5 | 0.35±0.04 | 0.35±0.09 | 0.17±0.06 |
|  |  | **/** | 8,93E-01 | 2,25E-01 |
|  | 1,8 | 0.33±0.04 | 0.39±0.07 | 0.11±0.02 |
|  |  | / | 2,73E-01 | 6,79E-02 |
|  | 2,1 | 0.32±0.07 | 0.43±0.15 | 0.11±0.03 |
|  |  | / | 2,85E-01 | 1,09E-01 |
| Complex | / | 0.73±0.01 | 0.67±0.01 | 0.33±0.02 |
|  |  | / | **5,73E-08** | **5,68E-36** |
| **Intensity coding** | | | | |
| **Category** | **/** | **CN** | **IC** | **AC** |
| Pure tones | / | 0.73±0.07 | 0.47±0.1 | 0.54±0.06 |
|  |  | **/** | **3,55E-02** | 5,51E-01 |
| Complex | / | 0.85±0.02 | 0.76±0.03 | 0.31±0.03 |
|  |  | / | **4,09E-02** | **6,55E-04** |

**Supplementary Table S1: Frequency and intensity statistics for the spatial code.**

Table summarizing the values and statistics of data plotted in **Fig. 4.** For each row, the top value is Mean±SEM for the region and the bottom value is the Wilcoxon rank-sum test between the region and the previous region (IC against CN, and AC against IC). Identity coding: Pure tones, N=8-3 sound pairs for 0,3-2,1 octave difference; Complex, N = 105 sound pairs. Intensity coding: Pure tones, N=14 sound pairs; Complex, N = 15 sound pairs. Significant differences are marked in bold.

| **Amplitude modulation coding** | | | | |
| --- | --- | --- | --- | --- |
| **Category** | **ΔOctaves** | **CN** | **IC** | **AC** |
| Periodic modulations | 1 | 0.98±0.01 | 0.88±0.02 | 0.79±0.02 |
|  |  | **/** | **1,53E-06** | **2,85E-03** |
|  | 2 | 0.97±0.01 | 0.71±0.03 | 0.59±0.02 |
|  |  | **/** | **1,28E-06** | **9,98E-04** |
|  | 3 | 0.95±0.01 | 0.56±0.05 | 0.47±0.03 |
|  |  | **/** | **2,67E-05** | 8,65E-02 |
|  | 4 | 0.93±0.01 | 0.51±0.08 | 0.33±0.06 |
|  |  | **/** | **7,76E-04** | 1,21E-01 |
|  | 5 | 0.91±0.03 | 0.39±0.12 | 0.25±0.11 |
|  |  | **/** | **1,17E-02** | 5,75E-01 |
| Periodic modulations against pure tones | / | 0.8±0.01 | 0.47±0.03 | 0.46±0.02 |
|  |  | / | **8,42E-08** | **3,41E-02** |
| Linear modulations | Up versus Down | 0.98±0.01 | 0.9±0.04 | 0.85±0.03 |
|  |  | / | **2,09E-02** | 1,55E-01 |
| Linear modulations against pure tones | Similar starting intensity | 0.87±0.04 | 0.74±0.06 | 0.86±0.05 |
|  |  | **/** | **2,62E-02** | 6,30E-02 |
|  | Opposite starting intensity | 0.89±0.03 | 0.81±0.04 | 0.73±0.05 |
|  |  | **/** | **1,42E-02** | 1,70E-01 |

**Supplementary Table S2: Amplitude modulation statistics for the spatial code.**

Table summarizing the values and statistics of data plotted in **Fig. 5**. For each row, the top value is Mean±SEM for the region and the bottom value is the Wilcoxon rank-sum test between the region and the previous region (IC against CN, and AC against IC). Significant differences are marked in bold. Periodic modulations N = 25 sound pairs for each difference; Periodic modulations against pure tones, N = 42 sound pairs; Linear modulations N = 13 sound pairs; Linear modulations against pure tones, N = 13 sound pairs;

| **Multi-frequency coding** | | | | |
| --- | --- | --- | --- | --- |
| **Category** | **/** | **CN** | **IC** | **AC** |
| Chords | / | 0.72±0.01 | 0.77±0.02 | 0.67±0.02 |
|  |  | **/** | **3,76E-10** | **1,90E-09** |
| Chords against pure tones | / | 0.89±0.01 | 0.87±0.01 | 0.74±0.03 |
|  |  | / | 4,79E-01 | **2,38E-05** |
| **Category** | **ΔOctaves** | **CN** | **IC** | **AC** |
| Noise bandwidth | 0,5 | 0.96±0.01 | 0.99±0.0 | 0.91±0.01 |
|  |  | **/** | **1,80E-02** | **1,80E-02** |
|  | 1 | 0.93±0.02 | 0.96±0.01 | 0.85±0.02 |
|  |  | **/** | 1,16E-01 | **2,77E-02** |
|  | 1,5 | 0.89±0.03 | 0.92±0.02 | 0.85±0.03 |
|  |  | **/** | **4,31E-02** | **4,31E-02** |
|  | 2 | 0.85±0.03 | 0.88±0.02 | 0.8±0.03 |
|  |  | / | 4,65E-01 | 6,79E-02 |
|  | 2,5 | 0.84±0.04 | 0.82±0.03 | 0.73±0.03 |
|  |  | / | 1,00E+00 | 1,09E-01 |
|  | 3 | 0.84±0.01 | 0.76±0.06 | 0.73±0.02 |
|  |  | / | 1,80E-01 | 6,55E-01 |
| **Category** | **Frequency content** | **CN** | **IC** | **AC** |
| Noise against pure frequencies | Broad | 0.76±0.02 | 0.55±0.04 | 0.46±0.03 |
|  |  | / | **9,18E-03** | **1,57E-02** |
|  | Ramps | 0.6±0.06 | 0.3±0.08 | 0.28±0.05 |
|  |  | / | **6,91E-03** | 3,86E-01 |

**Supplementary Table S3: Multi-frequency statistics for the spatial code.**

Table summarizing the values and statistics of data plotted in **Fig. 6**. For each row, the top value is Mean±SEM for the region and the bottom value is the Wilcoxon rank-sum test between the region and the previous region (IC against CN, and AC against IC). Significant differences are marked in bold. Chords N=195 sound pairs; Chords against pure tones, N =50 sound pairs; Noise bandwidth, N=8-3 sound pairs for 0,5-3 octave difference; Noise against pure tones, N =10 sound pairs for Broad, 8 sound pairs for Ramps;

| **Frequency modulation coding** | | | | |
| --- | --- | --- | --- | --- |
| **Category** | **/** | **CN** | **IC** | **AC** |
| Chirps direction | / | 0.98±0.01 | 0.91±0.02 | 0.65±0.06 |
|  |  | **/** | **4,31E-02** | **4,31E-02** |
| Chirps against pure tones | / | 0.95±0.01 | 0.68±0.04 | 0.61±0.05 |
|  |  | / | **5,06E-03** | 1,14E-01 |
| Complex direction | / | 0.98±0.0 | 0.96±0.01 | 0.84±0.02 |
|  |  | **/** | **3,56E-02** | **6,55E-04** |

**Supplementary Table S4: Frequency modulation statistics for the spatial code.**

Table summarizing the values and statistics of data plotted in **Fig. 7**. For each row, the top value is Mean±SEM for the region and the bottom value is the Wilcoxon rank-sum test between the region and the previous region (IC against CN, and AC against IC). Significant differences are marked in bold. Sweep direction N = 5 sound pairs; Sweep against pure tones, N = 10 sound pairs; Complex direction, N = 15 sound pairs;

| **Identity coding** | | | | |
| --- | --- | --- | --- | --- |
| **Category** | **ΔOctaves** | **CN** | **IC** | **AC** |
| Pure tones | 0,3 | 0,58±0,05 | 0,57±0,07 | 0,48±0,07 |
|  |  | / | 6,78E-01 | 4,41E-01 |
|  | 0,6 | 0,39±0,03 | 0,44±0,04 | 0,3±0,07 |
|  |  | **/** | 2,08E-01 | **3,57E-02** |
|  | 0,9 | 0,34±0,03 | 0,35±0,03 | 0,21±0,05 |
|  |  | **/** | 8,66E-01 | 9,10E-02 |
|  | 1,2 | 0,32±0,02 | 0,37±0,04 | 0,16±0,04 |
|  |  | **/** | 4,63E-01 | **2,77E-02** |
|  | 1,5 | 0,31±0,03 | 0,38±0,05 | 0,14±0,04 |
|  |  | **/** | 2,25E-01 | **4,31E-02** |
|  | 1,8 | 0,28±0,03 | 0,36±0,05 | 0,06±0,02 |
|  |  | / | 6,79E-02 | 6,79E-02 |
|  | 2,1 | 0,27±0,04 | 0,44±0,01 | 0,14±0,03 |
|  |  | / | 1,09E-01 | 1,09E-01 |
| Complex | / | 0,44±0,01 | 0,37±0,01 | 0,32±0,02 |
|  |  | / | **3,92E-23** | **3,36E-06** |
| **Intensity coding** | | | | |
| **Category** | **/** | **CN** | **IC** | **AC** |
| Pure tones | / | 0,67±0,06 | 0,48±0,09 | 0,49±0,06 |
|  |  | **/** | 5,55E-02 | 9,17E-01 |
| Complex | / | 0,73±0,02 | 0,65±0,02 | 0,32±0,03 |
|  |  | **/** | **2,68E-02** | **6,55E-04** |

**Supplementary Table S5: Frequency and intensity coding statistics for the spatio-temporal code.**

Table summarizing the values and statistics of data plotted in **Fig. S2**. For each row, the top value is Mean±SEM for the region and the bottom value is the Wilcoxon rank-sum test between the region and the previous region (IC against CN, and AC against IC). Identity coding: Pure tones, N = 8-3 sound pairs for 0,3-2,1 octave difference; Complex, N = 105 sound pairs. Intensity coding: Pure tones, N = 14 sound pairs; Complex, N = 15 sound pairs. Significant differences are marked in bold.

| **Amplitude modulation coding** | | | | |
| --- | --- | --- | --- | --- |
| **Category** | **ΔOctaves** | **CN** | **IC** | **AC** |
| Periodic modulations | 1 | 0,81±0,01 | 0,6±0,02 | 0,8±0,02 |
|  |  | / | **2,28E-07** | **1,70E-05** |
|  | 2 | 0,79±0,02 | 0,48±0,02 | 0,59±0,02 |
|  |  | **/** | **9,63E-07** | **1,22E-03** |
|  | 3 | 0,78±0,02 | 0,4±0,03 | 0,47±0,03 |
|  |  | **/** | **1,82E-05** | 1,10E-01 |
|  | 4 | 0,78±0,02 | 0,36±0,04 | 0,31±0,09 |
|  |  | **/** | **5,31E-04** | 2,78E-01 |
|  | 5 | 0,79±0,03 | 0,28±0,06 | 0,19±0,15 |
|  |  | **/** | **1,17E-02** | 8,89E-01 |
| Periodic modulations against pure tones | / | 0,8±0,01 | 0,47±0,03 | 0,46±0,02 |
|  |  | / | **1,77E-08** | 4,20E-01 |
| Linear modulations | Up versus Down | 0,8±0,04 | 0,62±0,07 | 0,71±0,06 |
|  |  | **/** | **2,08E-02** | 2,41E-01 |
| Linear modulations against pure tones | Similar starting intensity | 0,82±0,03 | 0,72±0,05 | 0,83±0,04 |
|  |  | **/** | **4,24E-02** | 7,92E-02 |
|  | Opposite starting intensity | 0,82±0,03 | 0,77±0,03 | 0,64±0,05 |
|  |  | **/** | 1,40E-01 | **3,31E-03** |

**Supplementary Table S6: Amplitude modulation statistics for the spatio-temporal code.**

Table summarizing the values and statistics of data plotted in **Fig. S3**. For each row, the top value is Mean±SEM for the region and the bottom value is the wilcoxon rank-sum test between the region and the previous region (IC against CN, and AC against IC). Significant differences are marked in bold. Periodic modulations N = 25 sound pairs for each difference; Periodic modulations against pure tones, N = 42 sound pairs; Linear modulations N = 13 sound pairs; Linear modulations against pure tones, N = 13 sound pairs;

| **Multi-frequency coding** | | | | |
| --- | --- | --- | --- | --- |
| **Category** | **/** | **CN** | **IC** | **AC** |
| Chords | / | 0,71±0,01 | 0,77±0,01 | 0,68±0,02 |
|  |  | **/** | **1,69E-09** | **5,03E-08** |
| Chords against pure tones | / | 0,89±0,01 | 0,87±0,01 | 0,74±0,03 |
|  |  | **/** | 1,63E-01 | **3,13E-05** |
| **Category** | **ΔOctaves** | **CN** | **IC** | **AC** |
| Noise bandwidth | 0,5 | 0,91±0,01 | 0,89±0,01 | 0,92±0,01 |
|  |  | / | **4,25E-02** | 1,28E-01 |
|  | 1 | 0,88±0,02 | 0,87±0,01 | 0,85±0,02 |
|  |  | **/** | 7,53E-01 | 3,45E-01 |
|  | 1,5 | 0,84±0,02 | 0,83±0,01 | 0,84±0,02 |
|  |  | **/** | 5,00E-01 | 8,93E-01 |
|  | 2 | 0,81±0,03 | 0,81±0,02 | 0,8±0,02 |
|  |  | / | 1,00E+00 | 7,15E-01 |
|  | 2,5 | 0,8±0,03 | 0,77±0,02 | 0,76±0,03 |
|  |  | / | 5,93E-01 | 2,85E-01 |
|  | 3 | 0,8±0,02 | 0,73±0,03 | 0,73±0,01 |
|  |  | / | 1,80E-01 | 6,55E-01 |
| **Category** | **Frequency content** | **CN** | **IC** | **AC** |
| Noise against pure frequencies | Broad | 0,69±0,02 | 0,5±0,02 | 0,42±0,01 |
|  |  | **/** | **9,82E-04** | **2,58E-02** |
|  | Ramps | 0,55±0,06 | 0,31±0,07 | 0,24±0,05 |
|  |  | / | **5,06E-03** | 1,39E-01 |

**Supplementary Table S7: Multi-frequency statistics for the spatial code.**

Table summarizing the values and statistics of data plotted in **Fig. S4**. For each row, the top value is Mean±SEM for the region and the bottom value is the wilcoxon rank-sum test between the region and the previous region (IC against CN, and AC against IC). Significant differences are marked in bold. Chords N = 195 sound pairs; Chords against pure tones, N = 50 sound pairs; Noise bandwidth, N = 8-3 sound pairs for 0,5-3 octave difference; Noise against pure tones, N = 10 sound pairs for Broad, 8 sound pairs for Ramps;

| **Frequency modulation coding** | | | | |
| --- | --- | --- | --- | --- |
| **Category** | **/** | **CN** | **IC** | **AC** |
| S5: Sweep direction | / | 0,4±0,1 | 0,27±0,08 | 0,31±0,11 |
|  |  | **/** | **4,31E-02** | 6,86E-01 |
| S5: Sweep against pure tones | / | 0,74±0,05 | 0,56±0,07 | 0,45±0,06 |
|  |  | / | **5,06E-03** | 1,14E-01 |
| S5: Complex direction | / | 0,74±0,02 | 0,57±0,02 | 0,78±0,02 |
|  |  | / | **6,55E-04** | **6,55E-04** |
| S6: Reconstruction Sound against Reconstruction Response (Spatial) | / | 0,57±0,01 | 0,41±0,01 | 0,38±0,01 |
|  |  | / | 6,79E-02 | 1,00E+00 |
| S6: Reconstruction Sound against Reconstruction Response (Spatio-temporal) | / | 0.82±0.01 | 0.72±0.01 | 0.51±0.01 |
|  |  | / | 6,79E-02 | 6,79E-02 |

**Supplementary Table S8: Frequency modulation statistics for the spatial code.**

Table summarizing the values and statistics of data plotted in **Fig. S5** and **S6** For each row, the top value is Mean±SEM for the region and the bottom value is the Wilcoxon rank-sum test between the region and the previous region (IC against CN, and AC against IC). Significant differences are marked in bold. Sweep direction N=5 sound pairs; Sweep against pure tones, N = 10 sound pairs; Complex direction, N = 15 sound pairs; Reconstruction (spatial), N = 4 sound pairs; reconstruction (spatio-temporal), N = 4 sound pairs;
