## Supplementary material for "Feature-dependent decorrelation of sound representations across the auditory pathway": Full manuscript including supplementray material

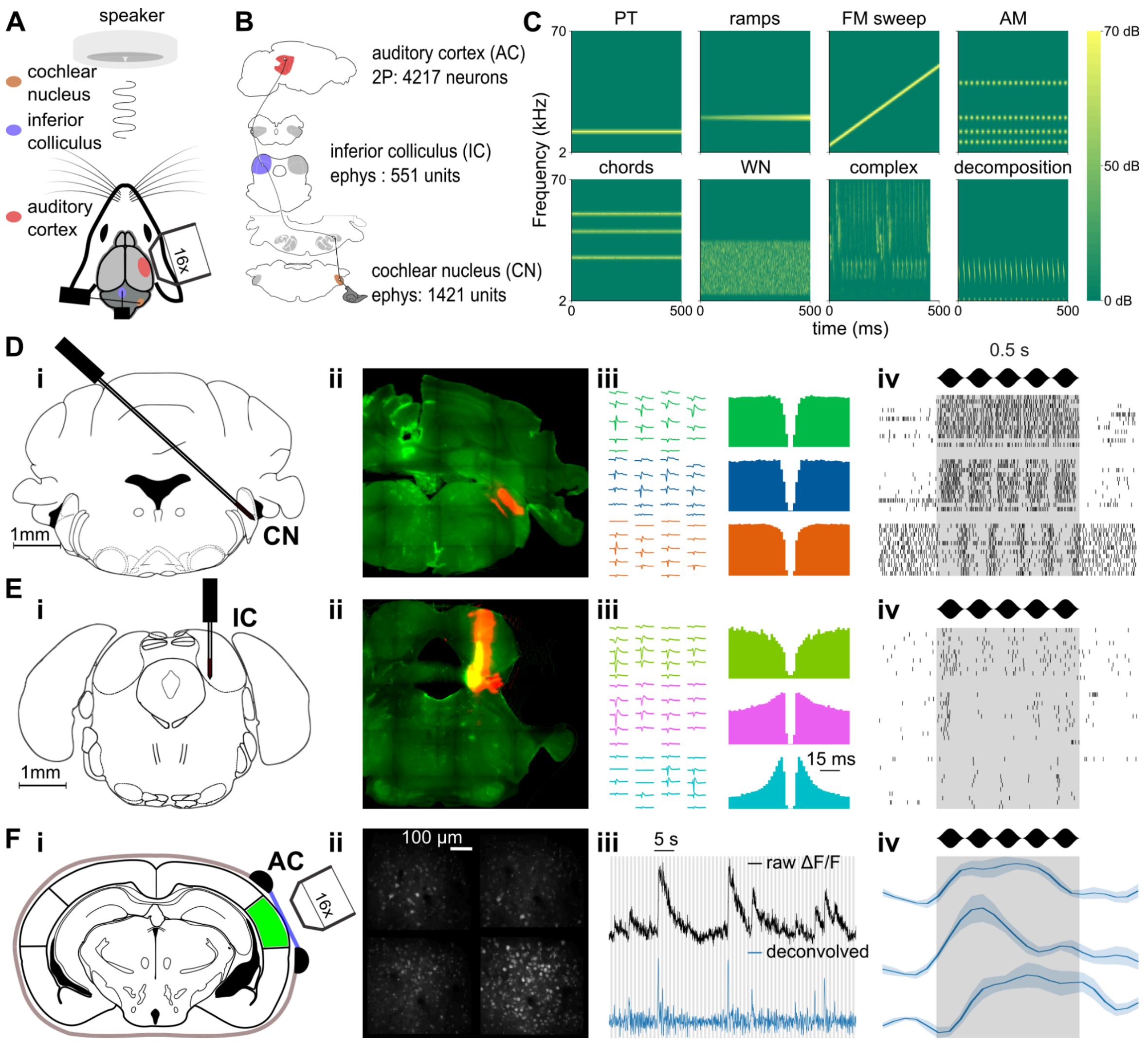


**Figure 1: Large-scale recordings of neuronal activity in head-fixed mice. A.** Schematic of the experimental setup. **B.** Schematic of the ascending auditory pathway. Each area where sound responses have been obtained via electrophysiology or 2-photon imaging is colored, with indication of the recording method and the number of single fibers/neurons. **C.** Sample spectrograms from each of the 8 categories contained in the 307 sounds stimulation set. **D-F.** Methodologies of data collection for each area. **D. i.** Schematic of the targeting of the cochlear nucleus using Neuropixels 1.0 probes **ii.** assessed via post-mortem histology. **iii.** Waveforms, auto-correlograms, **iv.** and raster plots of

responses to a 10 Hz AM of 3 example units. **E.** Same as **D** for inferior colliculus dataset collection. **F. i.** Schematic of the imaging of the auditory cortex **ii.** and example image of 4 planes recording. **iii.** Raw ΔF/F trace (black) and deconvolved trace (blue) in one example neuron from the imaging session in (**i, ii**). **iv.** Deconvolved response of 3 example neurons in response to a 10 Hz AM.

*General sharpening of sound representation tuning throughout the auditory system.* Plotting responses of individual neurons to sample sounds shows a large response diversity across sounds and neurons in the three considered stages of the auditory system (**Fig. 2**), reflecting the various neuronal subtypes and specializations present in these regions. Rich temporal variations of the firing rate are apparent mostly in electrophysiology recordings but also, at a lower resolution in calcium imaging recordings. This restricted temporal resolution is however not problematic because feature-specific tuning is defined as the specificity of the time-averaged firing rate of a neuron's response to particular stimuli, irrespective of response time course. Based on this definition, the representative examples shown in **Fig. 2** suggest that auditory cortex neurons have more specific responses, restricted to fewer sounds, as compared to subcortical neurons. To quantify this, we reasoned that more specifically tuned neurons lead, at the population level, to patterns of activated neurons that are more dissimilar across sounds. Hence, the overall statistics of single neuron tunings can be summarized in a population tuning measure that globally assesses how similar neural representations are across sounds.

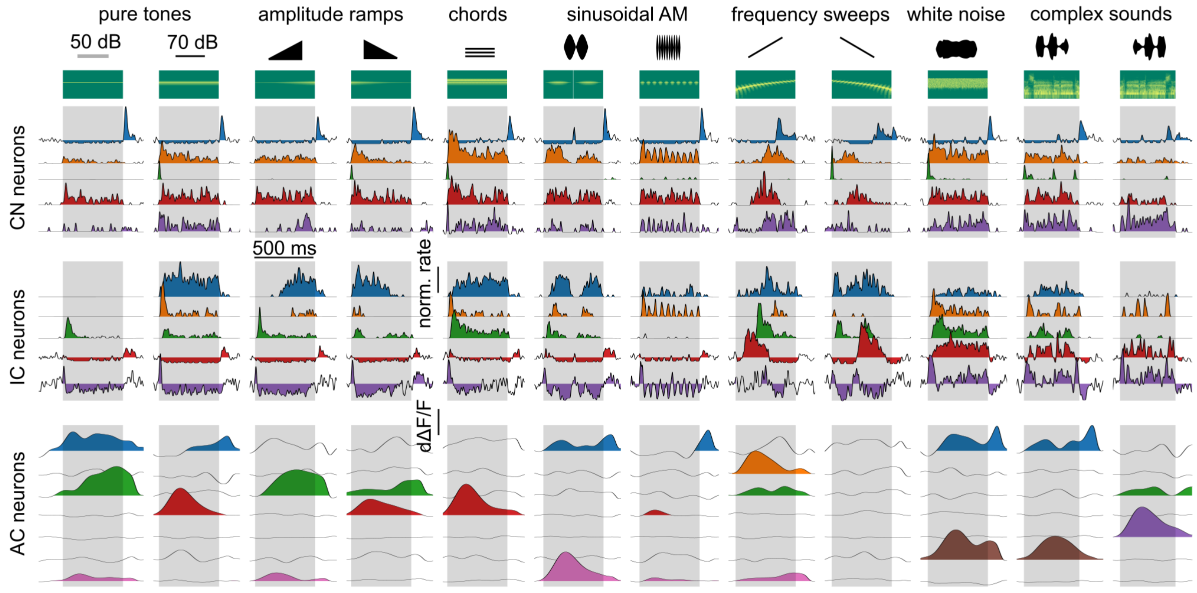


**Figure 2: Single cell sound response samples across the auditory system.** Trial-averaged responses of example neurons from cochlear nucleus (CN, 5 neurons), inferior colliculus (IC, 5 neurons), and auditory cortex (AC, 7 neurons) to 12 sounds with spectral content at 12 kHz (2 pure tones, 2 ramps, 1 chord, 2 AMs, 2 chirps, 1 WN and 2 complex, represented with their spectrograms). Sound presentation periods are shaded in gray.

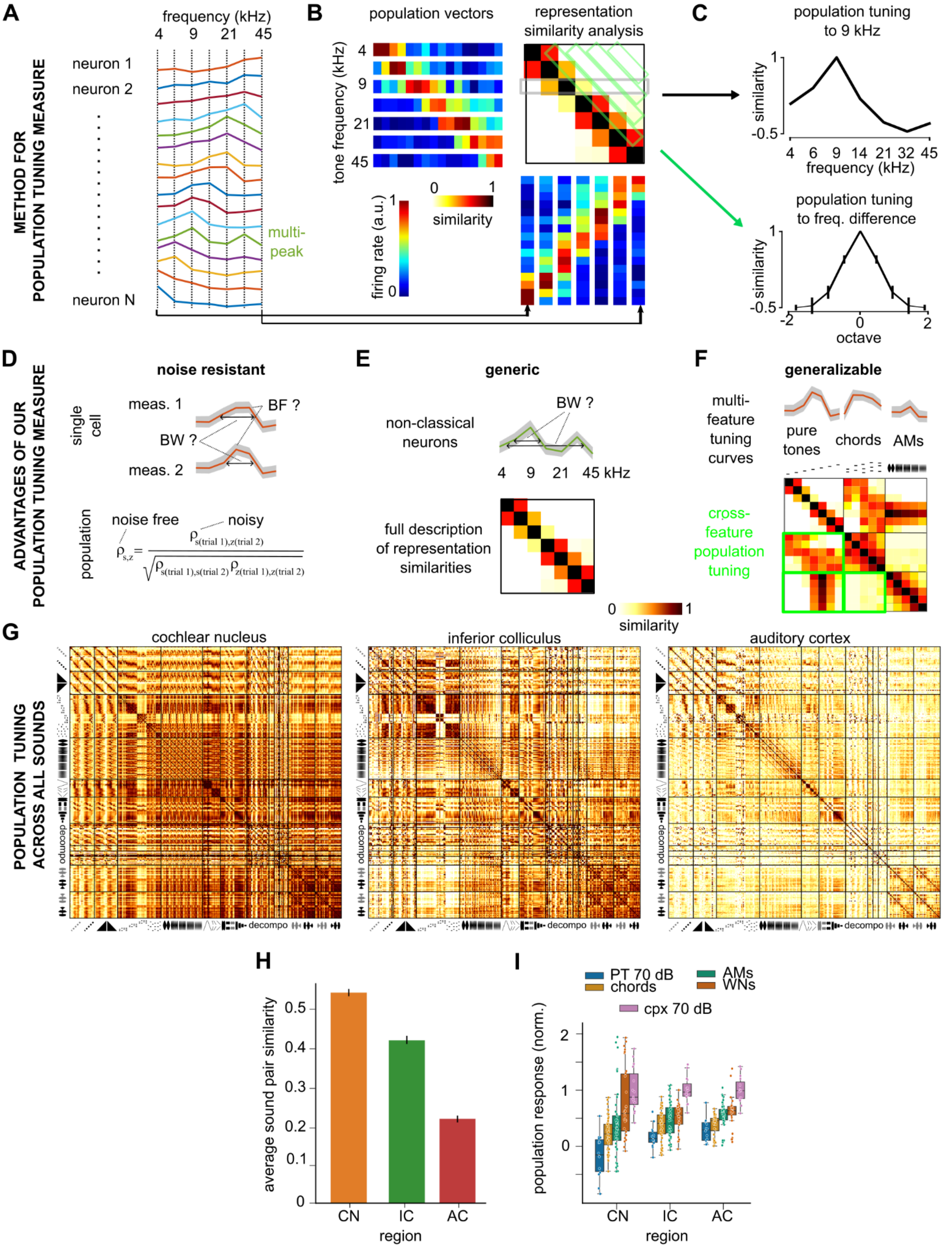


**Figure 3: Decorrelation of sound representations across the auditory system identified with a generic population tuning measure. A.** Synthetic tuning curves representing the time-averaged

firing rate responses of 16 modeled neurons to seven pure tones. **B**. Construction of the population similarity analysis matrix displaying the similarity of the population representations of the seven pure tones for the synthetic measures shown in **A**. **C**. For the same synthetic data, population tuning curves to 9kHz (*top*) and for different frequency ratios (measured in octaves) as extracted from the population similarity matrix in B. **D-F**. Illustration of three advantages of population tuning measures against single cell measurements. **D**. While single cell measurements of tuning properties such as best frequency (BF) and frequency tuning band-width (BW) are imprecise and biased because of response variability, the effect of noise on population tuning measures can be corrected. **E**. While single cell tuning measures often depend on a response model (e.g. single frequency peak), population tuning provides a generic description of the similarity relationships between all sounds (e.g. all pure tone frequencies). **F**. Population tuning is a unified measure that generalizes across any sound feature or any sound. **G.** Matrices of spatial representation similarity for all regions. **H.** Average spatial representation similarity between all pairs of sounds computed from **G** for all regions. (Mean±SEM: CN=0.53±1e-3, IC=0.41±1e-3, AC=0.21±1e-3. Two-sample Wilcoxon sign-rank test for paired distribution between pairs of sounds across regions: CN against IC, p<1e-63, IC against AC, p<1e-63). **I.** Average population response of neurons to distinct categories of sounds, after standardization of each neuron by its maximum response and of the population by the mean response to complex sounds at 70 dB SPL. CN = cochlear nucleus, IC = inferior colliculus, AC = auditory cortex.

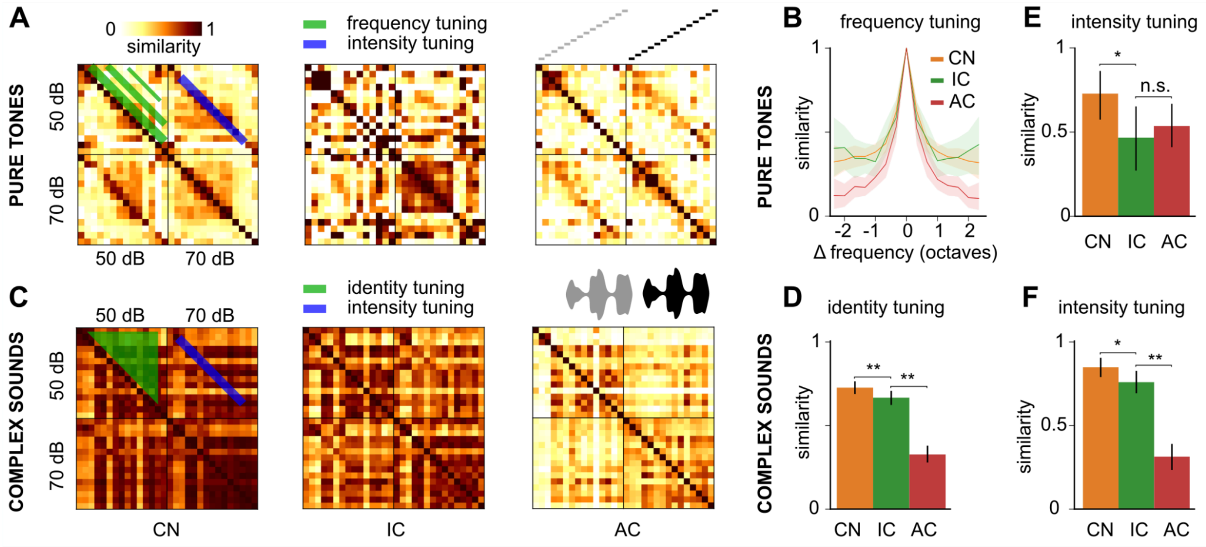


**Figure 4: Identity and intensity tuning improves differentially across the auditory system for simple and complex sounds. A.** Spatial representation similarity matrices for pure tones at 50- and 70-dB SPL. **B.** Evolution of spatial representation similarity between pure tones dependent on their frequency difference. **C.** Spatial representation similarity matrices for complex sounds at 50- and 70-dB SPL. **D.** Spatial representation similarity between different complex sounds with different identity (ie dolphin vs bird call) at the same average intensity. **E.** Spatial representation similarity between pure tones at the same frequency but at different intensities. **F.** Spatial representation similarity between identical complex sounds at different intensity. CN = cochlear nucleus, IC = inferior colliculus, AC = auditory cortex. Full statistics are provided in **Table S1**.

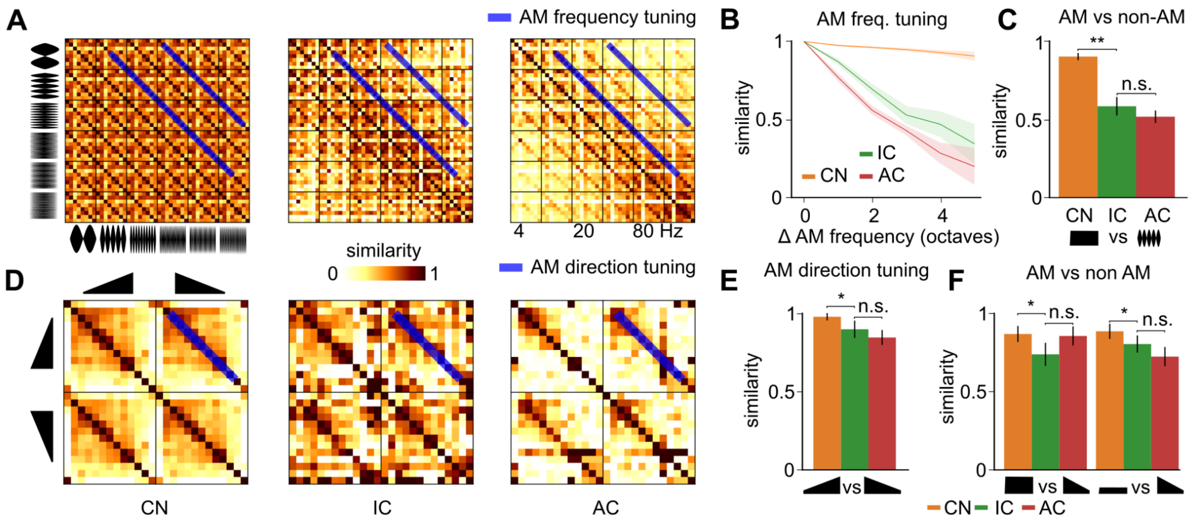


**Figure 5: Early emergence of amplitude modulation tuning. A.** Spatial representation similarity matrices for amplitude modulated (AM) sounds at 8 different carrier signals (pure tones and chords) and 6 modulation frequencies. **B.** Evolution of spatial similarity between AM sounds dependent on their modulation frequency difference. **C.** Similarity of the spatial representations of AM sounds and of the summed spatial representations of the pure tones corresponding to the carrier signal. **D.** Spatial representation similarity matrices for upward and downward linear intensity ramps (carrier signal = pure tone). **E.** Spatial representation similarity between upward and downward ramps at the same frequency. **F.** Spatial representation similarity between ramps and pure tones that have the same frequency and same start (l*eft*) or end (*right*) intensity as the ramp. CN = cochlear nucleus, IC = inferior colliculus, AC = auditory cortex. Full statistics are provided in **Table S2**.

*Cortical improvement of tuning to chords versus pure tones.* Natural sounds contain various frequency patterns which participate in the identification of the sound. We first studied harmonic and inharmonic combinations of pure frequencies that have a periodic or pseudo-periodic waveform. Our soundset includes a large number of frequency chords covering either low, middle, or high frequency ranges or the full frequency range, as well as a few harmonic chords (**Fig. 6A**). Tuning across different chords within the same frequency range (i.e. different combinations of the same subset of pure tones) was relatively stable across the auditory system with a small decrease in the inferior colliculus (**Fig. 6B** left, **Table S3**). Hence we did not observe a refinement in the representations of fine details in tone chords which is already accurate in the periphery, consistent with the lack of improvement of fine frequency tuning previously noted (**Fig. 4B**). Tone chord representations were slightly dissimilar to the summed representations of the corresponding pure tones in the cochlear nucleus. This similarity level remained stable in the colliculus and decreased significantly in the auditory cortex (**Fig. 6C, Table S3**). Hence, the main transformation of chord representations in the auditory system is their differentiation from pure tones in the cortex. Taking in account temporal information in the similarity analysis did not alter this conclusion (**Fig. S4**).

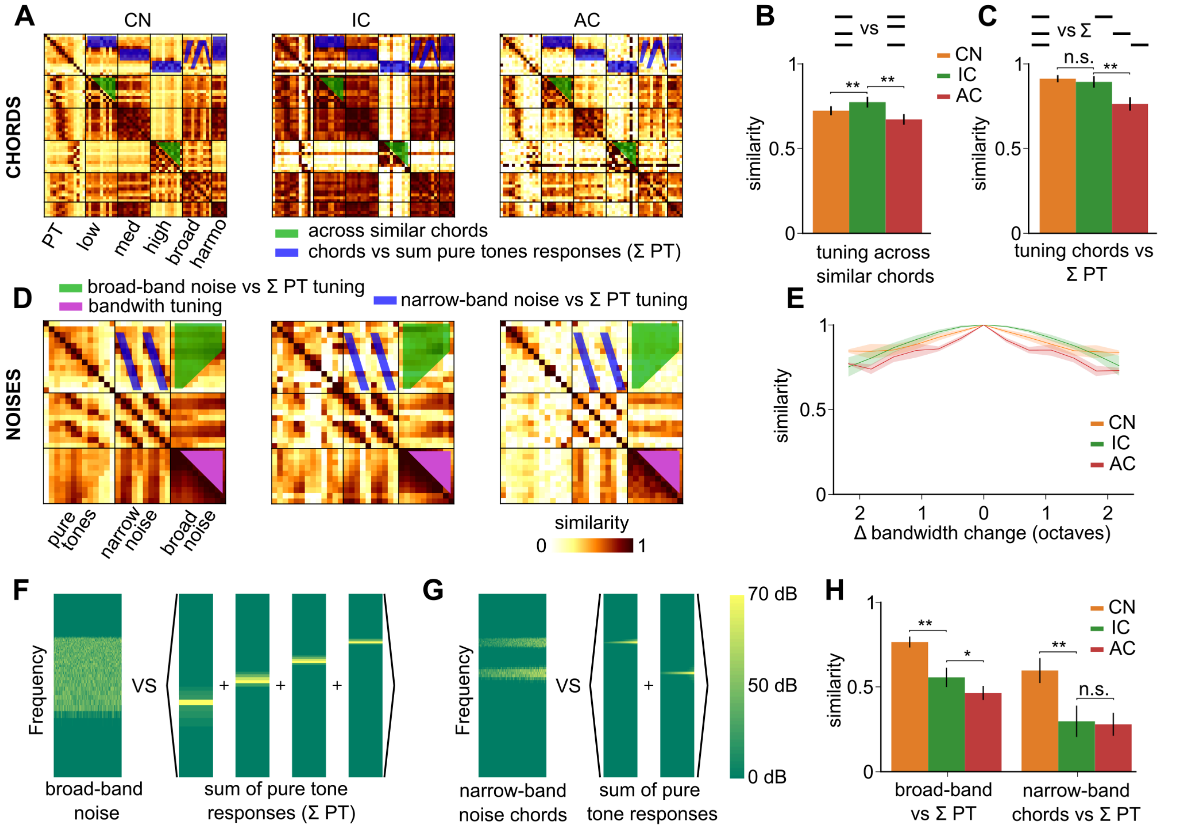


**Figure 6: Decorrelation of broad-band and multi-frequency sounds. A.** Spatial representation similarity matrices of pure tones at 70 dB SPL and their combination into various chords, organized based on the frequency range covered (low, mid, high, or full range) and on whether they are harmonic (harmo). **B.** Spatial representation similarity between chords built from the same pool of pure tones (*i.e.* between all pairs of chords built from low, mid, high frequency pure tones but not between a low frequency chord and a mid frequency chord). **C.** Similarity between spatial representations of chords and the summed representations of the pure tones that compose them. **D.** Spatial representation similarity matrices of pure tones, narrow- and broad-band noises. **E.** Mean spatial representation similarity between broadband noises dependent on their difference in bandwidth across recorded areas. **F.** Schematic of the reconstruction of a 4.8-28 kHz broadband noise using spectrograms of sounds used **G.** Schematic of the reconstruction of a 12+25 kHz narrowband noise

using spectrograms of sounds used. **H.** Similarity of spatial representations between broadband (left) or narrowband (right) noises and the summed response of pure tones included in their frequency

range. CN = cochlear nucleus, IC = inferior colliculus, AC = auditory cortex. Full statistics are provided in **Table S3**.

*Progressive refinement of tuning to broad- and narrow-band noises.* We next focused on broad-band frequency patterns which correspond to aperiodic, 'noise-like' sounds. To start exploring the tuning to broad-band sound parameters, we first quantified the similarity between population representations of filtered noises across different bandwidths, ranging from 3 kHz to 80 kHz, logarithmically centered around 10 kHz (**Fig. 6D**). Strikingly, we observed a weak tuning to noise bandwidth in all three considered brain regions (**Fig. 6E**). A slight improvement of tuning was observed from inferior colliculus to cortex. Broadband noise representations also differed from the summed representations of all sampled pure tones within their bandwidth, and more so in the inferior colliculus and cortex with the largest improvement occurring between cochlear nucleus and inferior colliculus (**Fig. 6F-H, Table S3**). Hence, like for amplitude modulations, most of the tuning to broadband sounds against pure tones is established in the inferior colliculus. Taking in account temporal information in the population tuning analysis did not substantially alter these observations (**Fig. S4**). Interestingly, broadband noises differ from pure tones in that their waveforms display large amplitude modulations of the envelope. Hence, the parallel improvement of tuning to broadband noise against pure tones and tuning to amplitude modulations (**Fig. 5**) suggests that part of the encoding of broadband noises in the colliculus is based on the detection amplitude modulations.

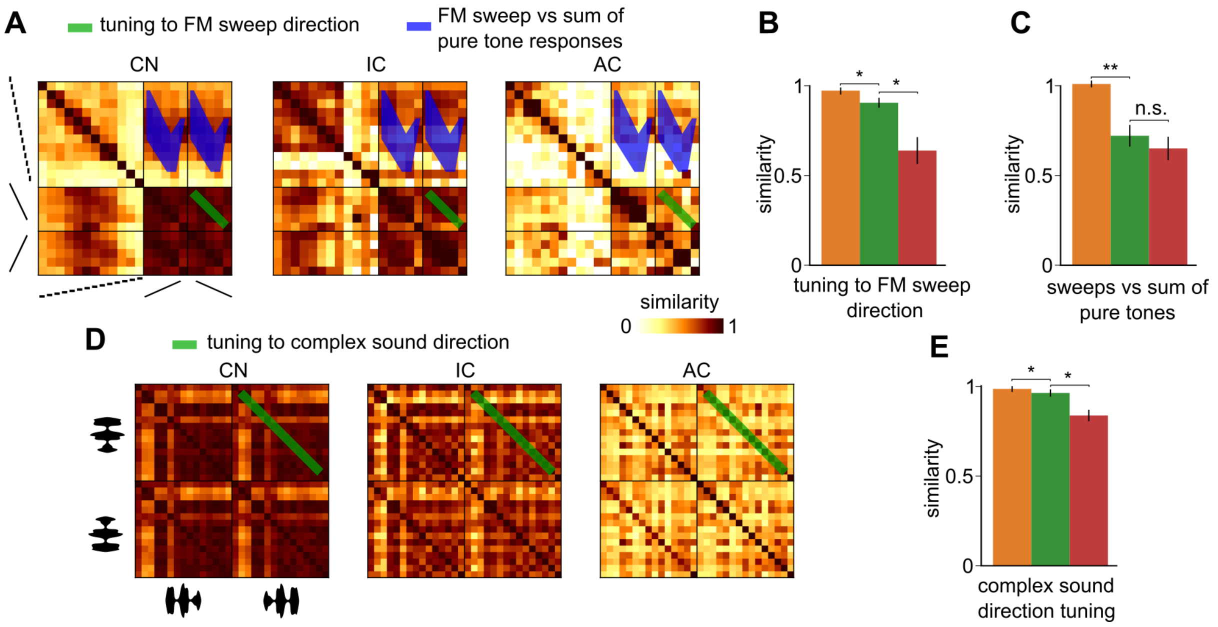


**Figure 7: Cortical decorrelation of sound direction. A.** Spatial representation similarity matrices of frequency modulated chirps at 70 dB SPL and pure tones at 70 dB SPL. **B.** Similarity of spatial representations between time-symmetric up- and down-frequency sweeps. **C.** Similarity of spatial representations of sweeps and of the summed spatial representations of pure tones traversed by the sweep. **D.** Spatial representation similarity matrices of forward and backward complex sounds at 70 dB SPL. **E.** Spatial representation similarity between forward and backward renditions of the same

complex sound. CN = cochlear nucleus, IC = inferior colliculus, AC = auditory cortex.

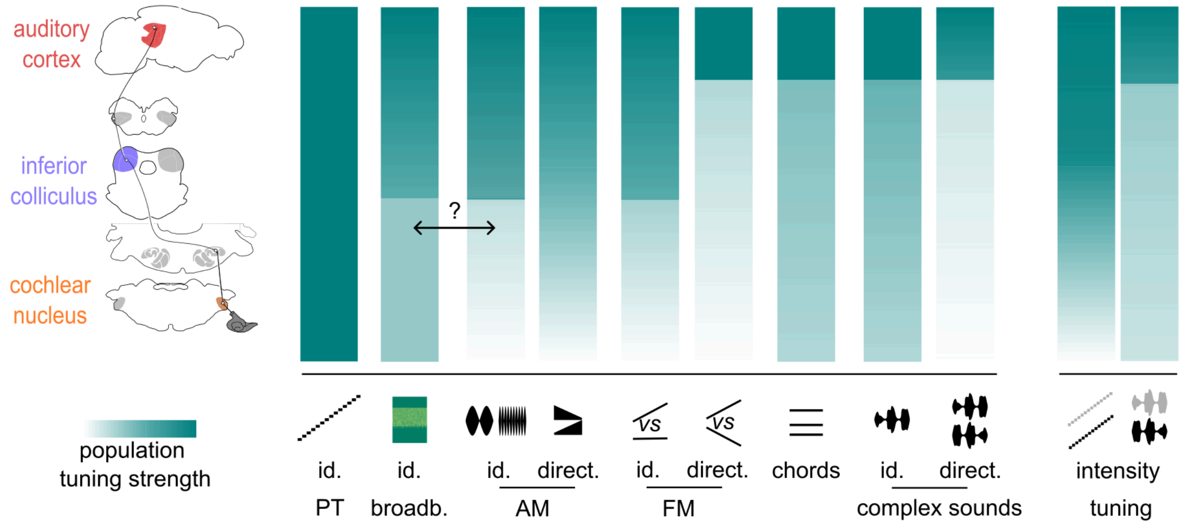


**Figure 8: Evolution of sound feature tuning along the auditory system.** Sketch representing population tuning strength across 4 different stages of the auditory (1 - similarity) for different

acoustic features from pure frequency tones to amplitude (AM) or frequency (FM) modulations to complex sounds. The double arrow indicates a possible link between tuning to AM and to noises. id. = identity ; direct. = direction ; freq. = frequency.

**Declaration of interests**

The authors declare no competing interests.

**Declaration of generative AI and AI-assisted technologies**

The authors did not use generative AI and AI-assisted technologies at any step of the study and in the writing process.

$$\rho_{\vec{\nu}_s \vec{\nu}_{s'}} \approx \frac{\frac{1}{R^2} \sum_{r,r'} \rho_{\vec{\nu}_{s,r} \vec{\nu}_{s',r'}}}{\sqrt{\frac{1}{R^2(1-R)^2} \left( \sum_{r \neq r'} \rho_{\vec{\nu}_{s,r} \vec{\nu}_{s,r'}} \right) \left( \sum_{r \neq r'} \rho_{\vec{\nu}_{s',r} \vec{\nu}_{s',r'}} \right)}}$$

in which r and r' are single trial indices and R is the total number of trials [19].

33. Kanold PO, Nelken I, Polley DB. Local versus global scales of organization in auditory cortex.

Trends Neurosci. 2014;37: 502–10. doi:S0166-2236(14)00100-3 [pii] 10.1016/j.tins.2014.06.003

### Supplementary figures

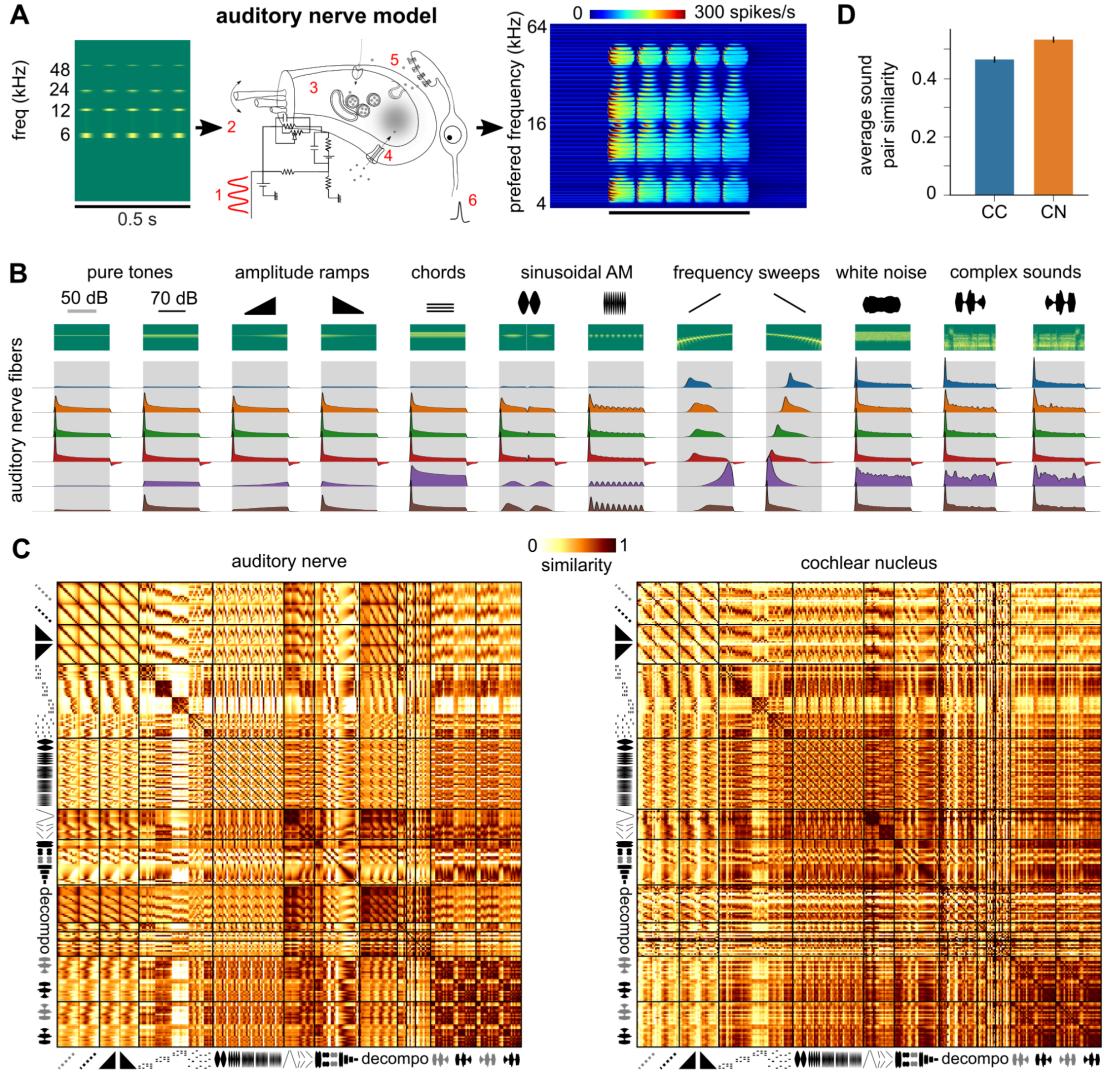


**Figure S1**: **Comparison of representations between cochlear nucleus and auditory nerve fiber model. A.** Sounds sampled at 400 kHz (represented here as a spectrogram) are fed into a detailed biophysical model of the cochlea and auditory nerve. The model simulates passive basilar membrane properties (1), stereocilia transduction (2, 3), calcium channel dynamics in hair cells (4) and synaptic release, as summarized in the Methods section. **B.** Responses of example neurons from the auditory nerve fiber model with spectral content at 12 kHz (2 pure tones, 2 ramps, 1 chord, 2 AMs, 2 chirps, 1 WN and 2 complex, represented with their spectrograms). Sound presentation periods are shaded in gray. **C.** Matrices of spatial representation similarity for the auditory nerve fiber model activity during sound presentation and for cochlear nucleus data as in **Fig. 3**. **D.** Average spatial representation similarity between all pairs of sounds computed from **C** for the auditory nerve fiber model and for cochlear nucleus data.

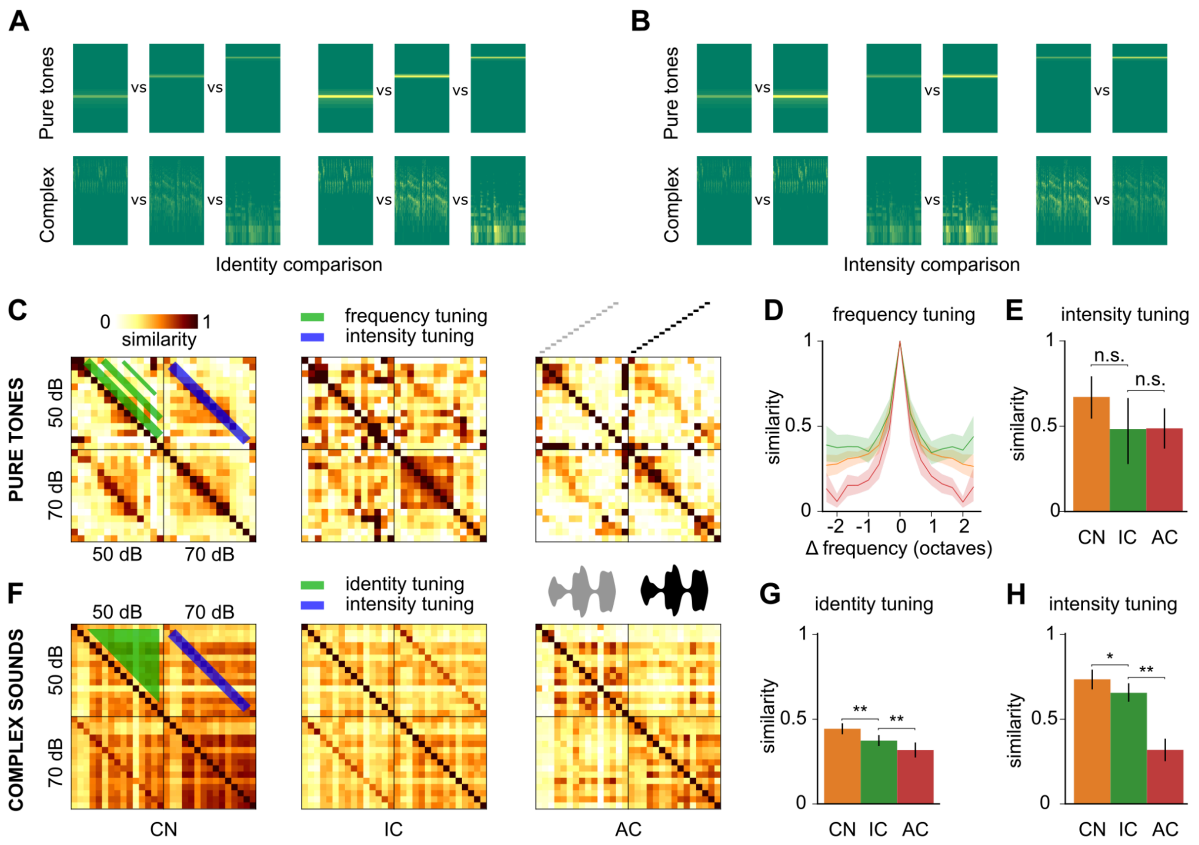


**Figure S2**: **Representation similarity of sound identity and intensity when including temporal information. A & B.** Sample spectrograms of the sounds compared in the population tuning analyses presented in **Fig. 4** and **S2C-H** separating identity and intensity comparisons. **C-H** Same measures of population tuning as in **Fig. 4A-F** but taking in account temporal information by comparing the concatenated population vector time series describing the full spatio-temporal population representation over the duration of the sound instead of using only the time-averaged activity. Temporal information does not change representation similarity for pure tones and pure tone intensity (**C-E**), but decreases similarity for complex sound identity and intensity (**F-H**). For identity, the result is expected as complex sounds include a large amount of temporal information. Complex sound decorrelation is therefore less pronounced for the spatio-temporal than for the time-averaged spatial representation (**Fig. 4E**), in line with the observation that a key computation of the auditory cortex is to encode temporal information into specific spatial patterns [19]. The improvement of intensity tuning for complex sounds (**H**) and not for pure tones after including temporal information was also observed previously [19]. CN = cochlear nucleus, IC = inferior colliculus, AC = auditory cortex. Full statistics are provided in **Table S5**.

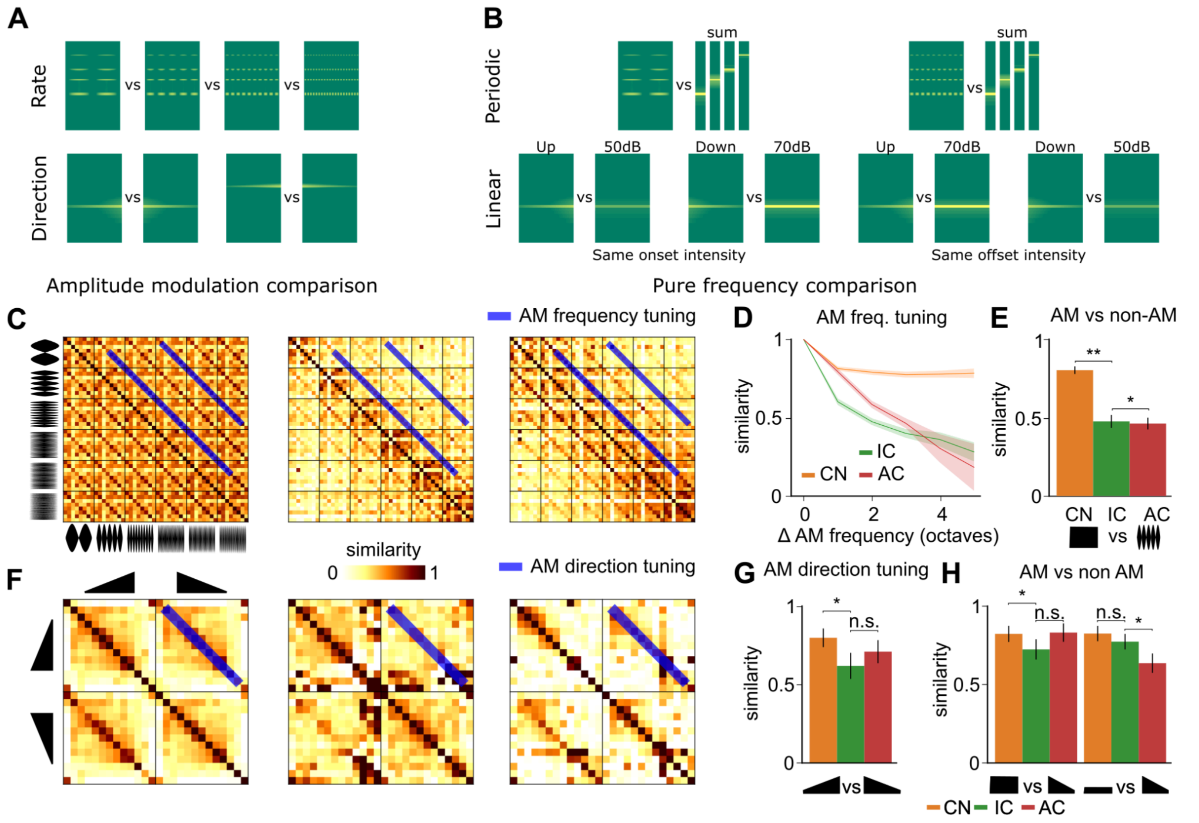


**Figure S3: Representation similarity of amplitude modulations when including temporal information. A.** Spectrogram of amplitude modulated sounds. **B.** Spectrogram summarizing the comparison between AM and non-AM activity. **C.** Spatio-temporal representation similarity matrices for 48 AM sounds at 8 carrier frequency contents and 6 modulation frequencies. **D.** Evolution of spatio-temporal similarity between AMs dependent on their modulation frequency difference. **E.** Spatio-temporal representation similarity between AM sounds and the non-modulated carrier signal. **F.** Spatio-temporal representation similarity matrices for upward and downward linear intensity ramps. **G.** Spatio-temporal similarity between upward and downward Ramps at the same frequency. **H.** Similarity between intensity ramps and pure tones at the same frequency and at the start (left) or end (right) intensity of the ramp. CN = cochlear nucleus, IC = inferior colliculus, AC = auditory cortex. Full statistics are provided in **Table S6**.

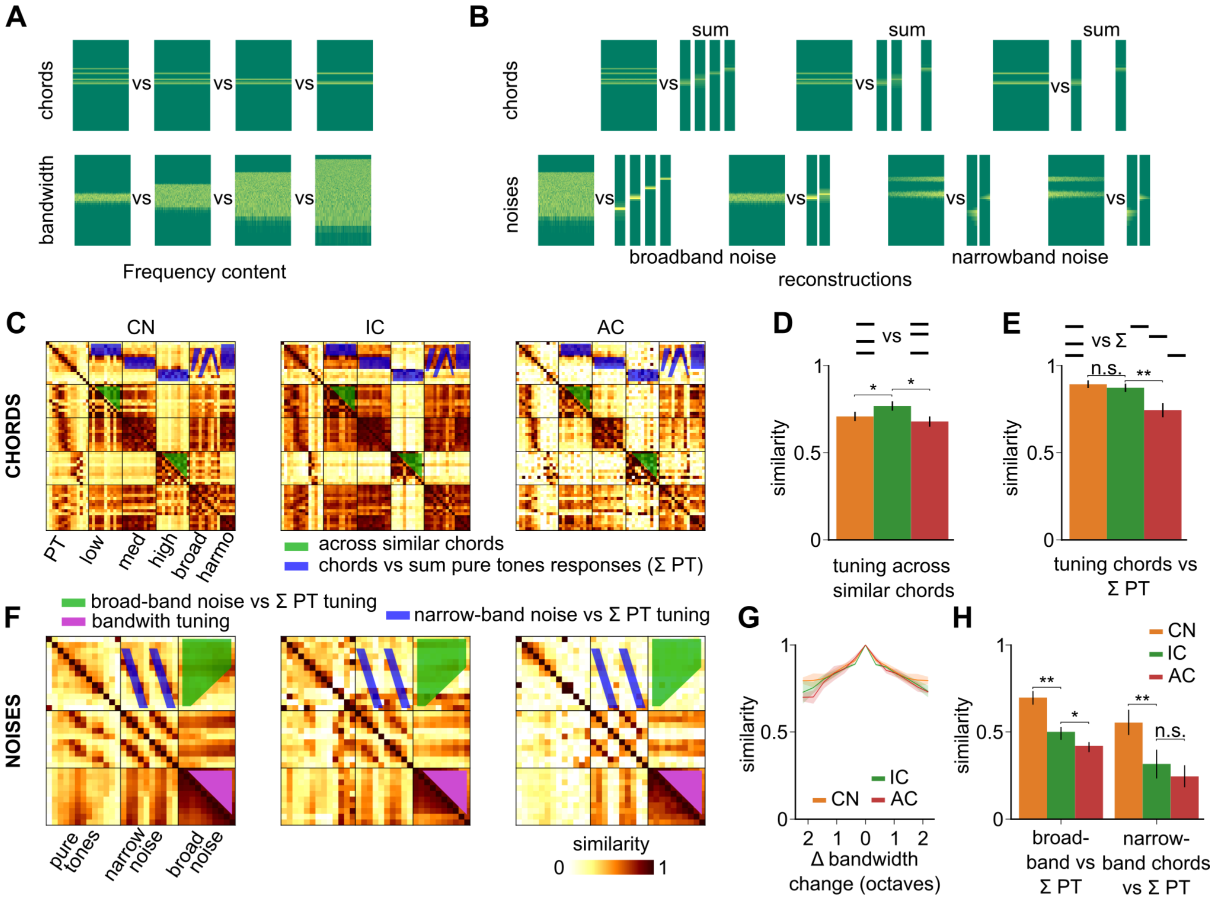


**Figure S4**: **Representation similarity of broad-band and multi-frequency sounds when including temporal information. A.** Spectrogram of different chords generated from the same pool of pure tones and of noises with different bandwidths. **B.** Spectrogram sketching the comparison between representation of 3 chords and the summed representations of the corresponding pure tones (top). Spectrogram sketching the comparison between representations of 4 noises and the summed representations of the pure tones included in their frequency range (bottom). **C.** Spatio-temporal representation similarity matrices of pure tones at 70 dB SPL and their summation into chords, organized based on their frequency content. **D.** Similarity between chords built from the same pool of pure tones. **E.** Similarity between spatio-temporal representations of chords and the summed representations of the pure tones that compose them. **F.** Spatio-temporal representation similarity matrices of pure tones, narrow- and broad-band noises. **G.** Spatio-temporal representation similarity between broadband noises dependent on their difference in bandwidth across recorded areas. **H.** Similarity of spatio-temporal representations between broadband (left) or narrowband (right) noises and the summed response of pure tones included in their frequency range. CN = cochlear nucleus, IC = inferior colliculus, AC = auditory cortex. Full statistics are provided in **Table S7**.

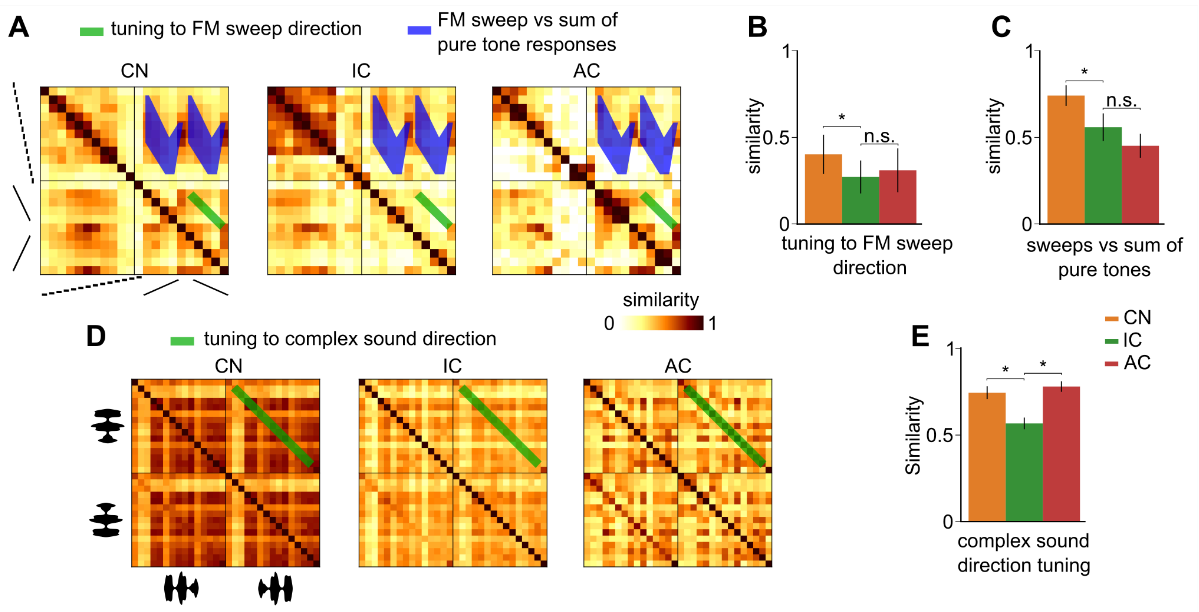


**Figure S5: Representation similarity for sound direction when including temporal information.** **A.** Representational similarity matrices of pure tones at 70 dB SPL and Chirps at 70 dB SPL that span the equivalent frequency range for the full spatio-temporal description of population activity in all regions. **B.** Similarity between same frequency up- and down-frequency modulated chirps for spatio-temporal population activity. **C.** Similarity between Chirps and the response reconstructed from pure tones traversed by the Chirp, for the spatial code. **D.** Representational similarity matrices of forward and backward Complex sounds at 70 dB SPL for spatio-temporal population activity in all regions. **E.** Similarity between the same complex sounds in both temporal directions for spatio-temporal population activity. CN = cochlear nucleus, IC = inferior colliculus, AC = auditory cortex. CN = cochlear nucleus, IC = inferior colliculus, AC = auditory cortex. Full statistics are provided in **Table S8**.

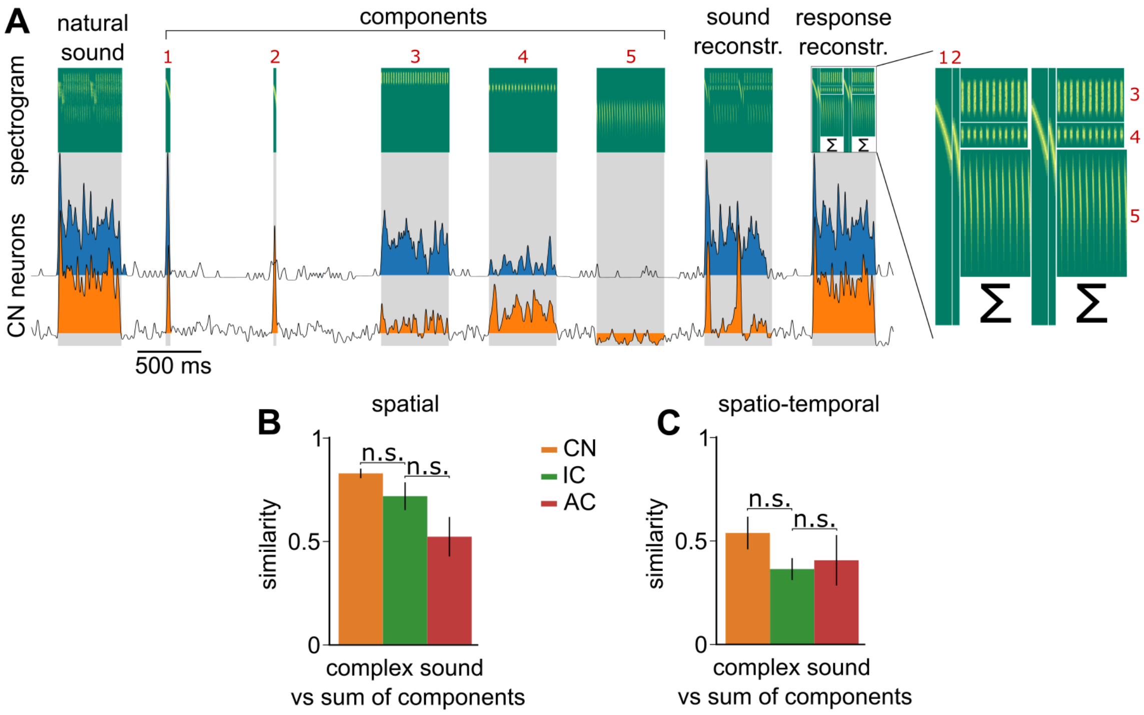


**Figure S6**: **Specificity of complex sound representations against the sum of their decomposition. A.** Schematic of the reconstruction process of sounds as a sum of individual components (top) and responses of 2 example neurons from the cochlear nucleus to the example complex sound, its components, its reconstruction, and its reconstructed response). **B.** Similarity between the responses to reconstructed complex sounds and reconstructed response from responses to components of the complex sound for 3 complex sounds for both codes. **C**. same as **B** but for the full spatio-temporal population activity. CN = cochlear nucleus, IC = inferior colliculus, AC = auditory cortex. Full statistics are provided in **Table S8**.
